# The zinc cluster transcription factor Rha1 is a positive filamentation regulator in *Candida albicans*

**DOI:** 10.1101/2020.01.21.901744

**Authors:** Raha Parvizi Omran, Chris Law, Vanessa Dumeaux, Joachim Morschhäuser, Malcolm Whiteway

## Abstract

Zinc cluster transcription factors are essential fungal specific regulators of gene expression. In the dimorphic pathogen *Candida albicans,* they control processes ranging from metabolism and stress adaptation to mating, virulence, and antifungal resistance. Here, we have identified the gene *CaORF19.1604* as encoding a zinc cluster transcription factor that acts as a regulator of filament development. Hyperactivation of *CaORF19.1604*, which we have named *RHA1* for <u>R</u>egulator of <u>H</u>yphal <u>A</u>ctivity, leads to a wrinkled colony morphology under non-hyphal growth conditions, to pseudohyphal growth and filament formation, to invasiveness and enhanced biofilm formation.  Cells with activated Rha1 are sensitive to cell wall modifying agents such as Congo red and the echinocandin drug caspofungin but show normal sensitivity to fluconazole. RNA-sequencing-based transcriptional profiling of the activated Rha1 strain reveals the up-regulation of genes for core filamentation and cell-wall-adhesion-related proteins such as Als1, Als3, Ece1, and Hwp1. Upregulation is also seen for the genes for the hyphal-inducing transcription factors Brg1 and Ume6 and genes encoding several enzymes involved in arginine metabolism, while downregulation is seen for the hyphal repressor Nrg1. The deletion of *BRG1* blocks the filamentation caused by activated Rha1, while null mutants of *UME6* result in a partial block. Deletion of *RHA1* can partially reduce healthy hyphal development triggered by environmental conditions such as Spider medium or serum at 37°C.

In contrast to the limited effect of either single mutant, the double *rha1 ume6* deletion strain is totally defective in both serum and Spider medium stimulated hyphal development. While the loss of Brg1 function blocks serum-stimulated hyphal development, this block can be significantly bypassed by Rha1 hyperactivity, and the combination of Rha1 hyperactivity and serum addition can generate significant polarization in even *brg1 ume6* double mutants. Our results thus suggest that in response to external signals, Rha1 functions to facilitate the switch from an Nrg1 controlled yeast state to a Brg1/Ume6 regulated hyphal state.

**Author Summary:** *Candida albicans* is the predominant human fungal pathogen, generating a mortality rate of 40% in systemically infected patients. The ability of *Candida albicans* to change its morphology is a determinant of its tissue penetration and invasion in response to variant host-related stimuli. The regulatory mechanism for filamentation includes a complex network of transcription factors that play roles in regulating hyphae associated genes. We identify here a new regulator of filamentation from the zinc cluster transcription factor family. We present evidence suggesting that this transcription factor assists the Nrg1/Brg1 switch regulating hyphal development.

## Introduction

Transcriptional control of cellular processes is critical for normal functioning of the medically important opportunistic fungal pathogen, *Candida albicans*, Central to this regulation are the DNA-binding transcription factors (TFs) that generally associate with target sequences in the promoters of regulated genes. The bound factors serve to activate or repress transcription in response to signals that typically represent either internal cellular states or external conditions (1, 2). One of the main classes of *C. albicans* transcription factors are the zinc cluster proteins, named because of a cysteine-rich region that coordinates a zinc atom as part of the DNA binding domain of the protein (3). Such zinc cluster transcription factors (ZCTFs) are found only in fungi and amoebae, but related zinc finger proteins serve as transcription factors throughout the eukaryotes (3, 4). Other *C. albicans* transcription factor classes are categorized into the basic helix-loop-helix (bHLH) class, zinc finger TFs, homeobox family TFs, and leucine zipper TFs (5, 6). Many studies have been conducted to characterize the ZCTF function; a powerful tool in these studies has been the use of activated versions of the ZCTFs. For example, screening by Schillig *et al*.(7) of a comprehensive collection of Zn(II)2 Cys6 gain-of-function mutants identified regulatory genes controlling fluconazole resistance, including Mrr2 (multidrug resistance regulator) that regulates the multidrug efflux pump Cdr1. Tebung *et al.* (8) used ZCTF activation during the characterization of a case of rewiring between purine and pyrimidine metabolism in the ascomycetes, and recently they found a conserved role for Put3 in regulating proline catabolism in both *C. albicans* and *S. cerevisiae* (9). These studies led us to investigate ZCFs further to uncover the transcriptional regulatory circuits involved in pathogenicity critical processes such as hyphal growth.

An essential characteristic of *C. albicans* that has been critical for its success as a commensal and as an opportunistic pathogen is its ability to grow in different morphological forms. The two most common cellular forms are the yeast form, where cells grow by budding as individual, rounded cells, and the hyphal form, where cells grow as extended, branching filaments, with individual cells delineated by septa (10). *C. albicans* cells use the filament form to escape from human macrophages and to invade into the deeper tissue during infection (11–13). Transcriptional control is essential for the ability of cells to make the switch between the yeast and filamentous forms. Several transcription factors have been identified that function in this switch. These include Efg1, a target of the Ras1/cAMP pathway critical for hyphal development (14), Cph1, a target of the MAP kinase pathway (15), Tec1 (16). They also include transcriptional repressors such as Tup1 (17), and Nrg1 (18).

The molecular mechanism of the hyphal initiation step suggests that environmental stimuli such as serum, Spider medium, and GlcNac, serve to trigger a switch from a yeast state, where the Nrg1 repressor blocks activity of hyphal-associated genes (HAGs), to a hyphal state, where the Brg1 transcription factor and the associating Hda1 histone deacetylase remodel the chromatin state of the HAG promoters. This remodeling leads to the activation of the hyphal program (19). Thus, hyphal development involves downregulation of transcription factor Nrg1 by the cAMP-dependent PKA pathway, together with upregulation of Brg1 and Ume6 (18), and It has been proposed that the chromatin state of the promoters of crucial genes represents a central regulator of transcriptional control of the transition between the yeast and hyphal states (20). This circuit regulating chromatin structure required for hyphal initiation and extension involves the GATA family member transcription factor Brg1 (19) and Ume6, a zinc cluster family member (21, 22). Brg1 expression is repressed by Nrg1 (23), and Brg1 and Nrg1 compete to control the chromatin state.

Our work identifies a new transcription factor involved in the hyphal transition. The zinc cluster transcription factor Rha1 serves as a critical regulator of the Nrg1/Brg1 switch; activated Rha1 can trigger filamentous growth in the absence of external signals, and in the presence of external signals like serum, stimulation can bypass the need for Brg1. Loss of Rha1 function leads to reduced ability to generate hyphal growth in the presence of external signals, and when coupled with loss of Ume6 function, creates completely non-hyphal cells. This establishes Rha1 as a new ZCTF functioning in the hyphal control circuitry of the human pathogen *C. albicans*.

## Results

### Overexpression of Rha1 triggers *C. albicans* filamentation

We screened a library of *C. albicans* strains containing overexpressed and activated zinc cluster transcription factors (RW.ERROR - Unable to find reference:42), and identified Orf19.1604 as generating an abnormal colony morphology; colonies of cells over-expressing Orf19.1604 fused to a mutant Gal4 activation domain were more wrinkled and crenulated than the smooth colonies of the control strain SC5314 (Figure 1A). The activated strains were also invasive and resistant to washing from the surface of YPD medium plates after growth at 30°C. In contrast, cells of the control strain SC5314 were washed more easily from the surface of the agar (Figure 1B). When grown in liquid YPD medium at 30°C, strains containing activated Orf19.1604 were highly flocculant (Figure 1C), and assessment of the cellular morphology showed a high frequency of filamentous cells exhibiting primarily pseudohyphal characteristics (Figure 1D). As a consequence of this role in filamentation, we named *ORF19.1604 RHA1*, for the <u>R</u>egulator of <u>H</u>yphal <u>A</u>ctivity. Finally, we observed enhanced biofilm formation in the *RHA1* activated strain compared to strain SC5314 (Figure 1E).

**Figure 1.**
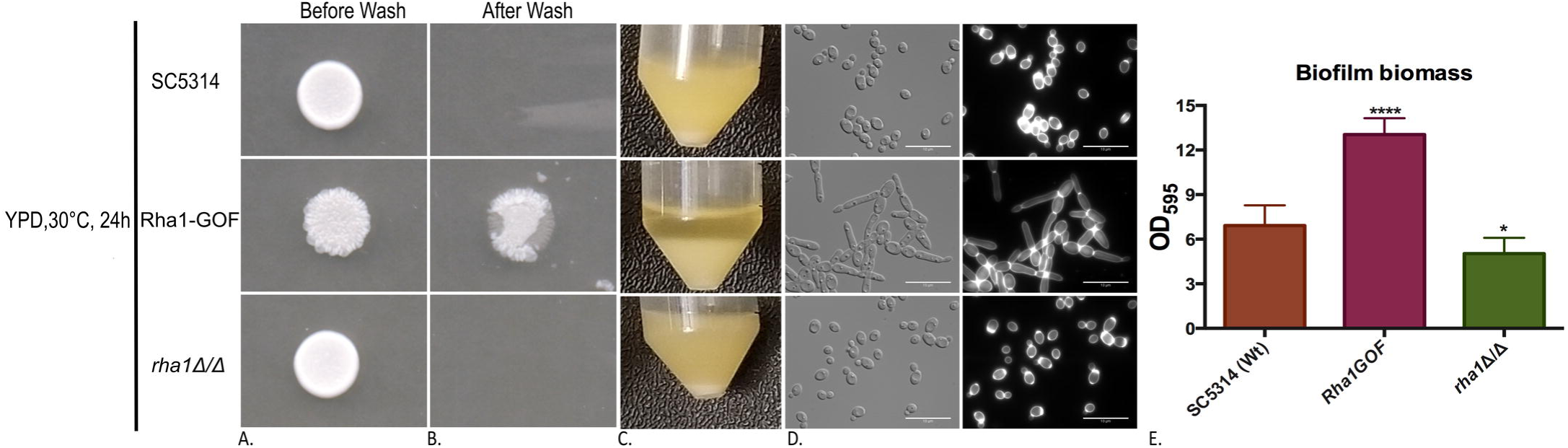
Morphological and biofilm results of the Sc5314 (WT), Rha1-GOF, and rha1 mutant. A) Sc5314 (Wild-type), Rha1-GOF, and *rha1 Δ/Δ* strains were spotted on the solid non-filament-inducing medium YPD and grown for 24hrs before washing with a stream of water for 15 seconds. The Rha1-GOF strain was invasive and resistant to washing. B) The indicated strains (Sc5314 (Wild-type), Rha1-GOF, and *rha1 Δ/Δ)* were grown in liquid YPD medium at 30 degrees. C) The cellular morphology of the indicated strains is displayed after growth overnight at 30 degrees in liquid YPD medium. Cells were washed twice with 1xPBS, stained with CFW, and visualized by DIC optics (Bar,10 µm). (E) A biofilm assay was run the indicated strains in Spider medium at 37°C after 24hrs. The experiments were assayed in triplicate. *t*-test, \**P* < 0.05, \*\**P* < 0.01, \*\*\**P* < 0.001

### Rha1 orthologs are limited to the CUG clade

In *C. albicans*, *RHA1* (C2_09460C_A) encodes a 989 amino acid protein with a zinc cluster type DNA-binding domain at the N terminal that shows limited sequence similarity to that of Lys14 in *Saccharomyces cerevisiae*, although Rha1 is unlike ScLys14 in the rest of the protein. In order to identify orthologs of *C. albicans* Rha1, the protein sequence from the Candida Genome Database (CGD) (http://www.candidagenome.org/) was compared across the ascomycetes using Blastp searches of the multiple fungal genome database (https://www.yeastgenome.org/blast-fungal). Convincing orthologs with sequence similarity outside of the DNA binding region of Rha1 are limited to CTG clade-specific species. They are not present in several phylogenetically related non-pathogenic species, including *Saccharomyces cerevisiae*, *S. paradoxus*, *S. mikatae*, *S. bayanus*, *S. castelli, and K. lactis* (Figure 2).

**Figure 2.**
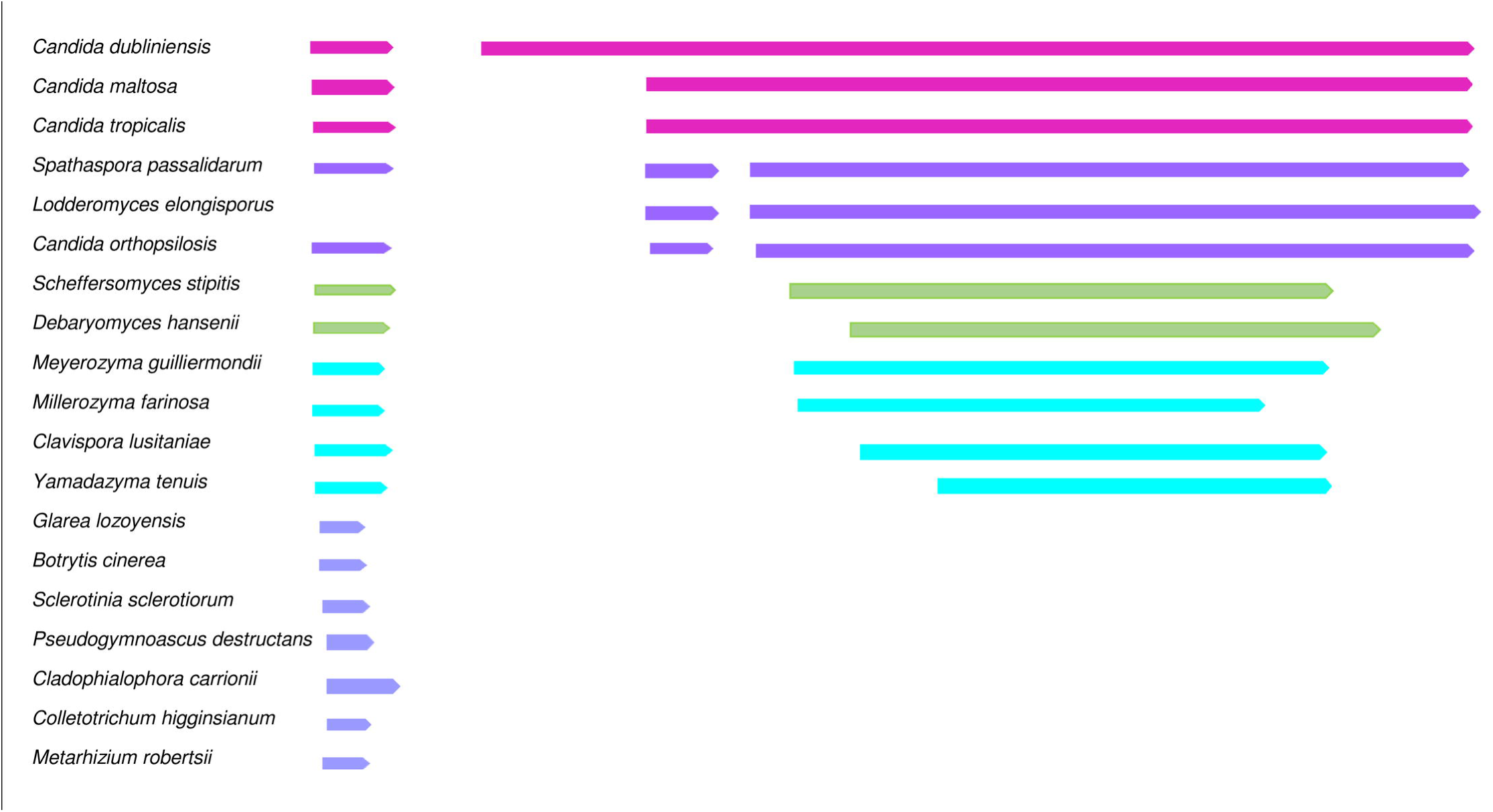
Blastp alignment of Rha1 through the Saccharomyces Genome Database (SGD). Protein sequence alignment of Rha1 with 964 amino acids versus different genome databases available in SGD. The multiple fungal sequence alignments outside the zinc cluster conserved motif establishes there is no protein sequence similarity of Rha1 with the non-CUG clade species.

### Activated Rha1 influences cell wall phenotypes

We examined the sensitivity of strains containing activated Rha1 to a variety of antifungal and cell wall stress drugs. Hyperactivity of Rha1 increases the sensitivity of cells to the echinocandin caspofungin and to Congo Red (Figure 3), compounds that affect the formation of the *C. albicans* cell wall through different mechanisms. By contrast, response to the antifungal drug fluconazole, which targets ergosterol biosynthesis and thus influences the *C. albicans* cell membrane, is not modified in strains carrying activated Rha1 (Figure 3). As well, the activated allele does not enhance sensitivity to the DNA damaging agent MMS 0.02%, the oxidative stress-inducing agent hydrogen peroxide (H_2_O_2_) 5mM, the salt stressors, 100mM Fe (II)SO_4_,20 mM Fe (III)Cl, and 5mM CuSO_4_, and the osmotic stressor 1M sorbitol 0.4m Cacl_2_, 0.15mM Menadione, HU 38mM, pH 10, Glycerol 250mM Hygromycin B 100µg/ml (Figure S1).

**Figure 3.**
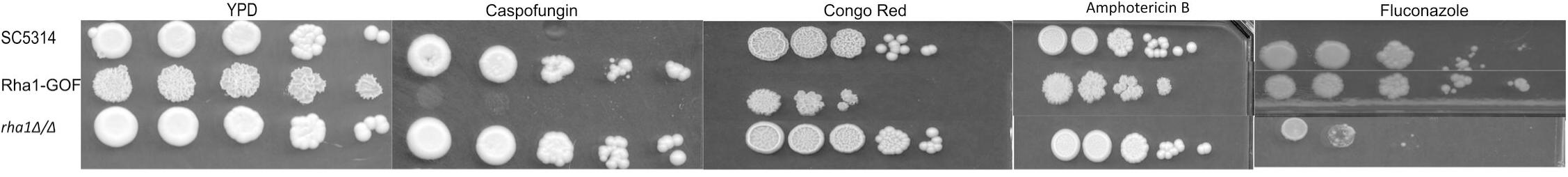
Response to cell wall stress and variable antifungal drugs of strains Sc5314, Rha1-GOF and rha1ΔΔ. Overnight cultures of the represented strains were washed twice in 1x PBS, then five microlitres of a 10-fold dilution were spotted under the indicated conditions for 48hrs at 30 degrees. Rha1-GOF is normally resistant to fluconazole and sensitive to the cell wall stresses.

### Activated Rha1 modulates gene expression

We performed RNA sequencing of duplicated samples to determine the transcriptional profile of cells carrying the activated *RHA1* allele compared to the wild type strain SC5314 cultured under the non-hyphae-inducing conditions of YPD medium at 30°C.  As shown in figure 4, three notable classes of genes have been up-regulated above 3-fold, with a P-value <0.03, compared to the reference strain SC5314. The first class of genes encodes classic cell wall adhesions such as Als1, Als3, and core filament-specific proteins such as Hwp1, Ece1, Ihd1, and Rbt1, consistent with the observed phenotype of filamentous cells (24). A second class consists of genes encoding transcription factors that themselves regulate hyphal development, specifically Brg1 and Ume6. The third notable collection of significantly upregulated genes encodes arginine metabolism enzymes such as Arg1, Arg3, Arg4, and the arginine permease Can2. As well, many genes appear downregulated in cells with hyperactive Rha1. Notable among these is gene for the hyphal repressor Nrg1, which is repressed 2.5-fold in these cells.

**Figure 4.**
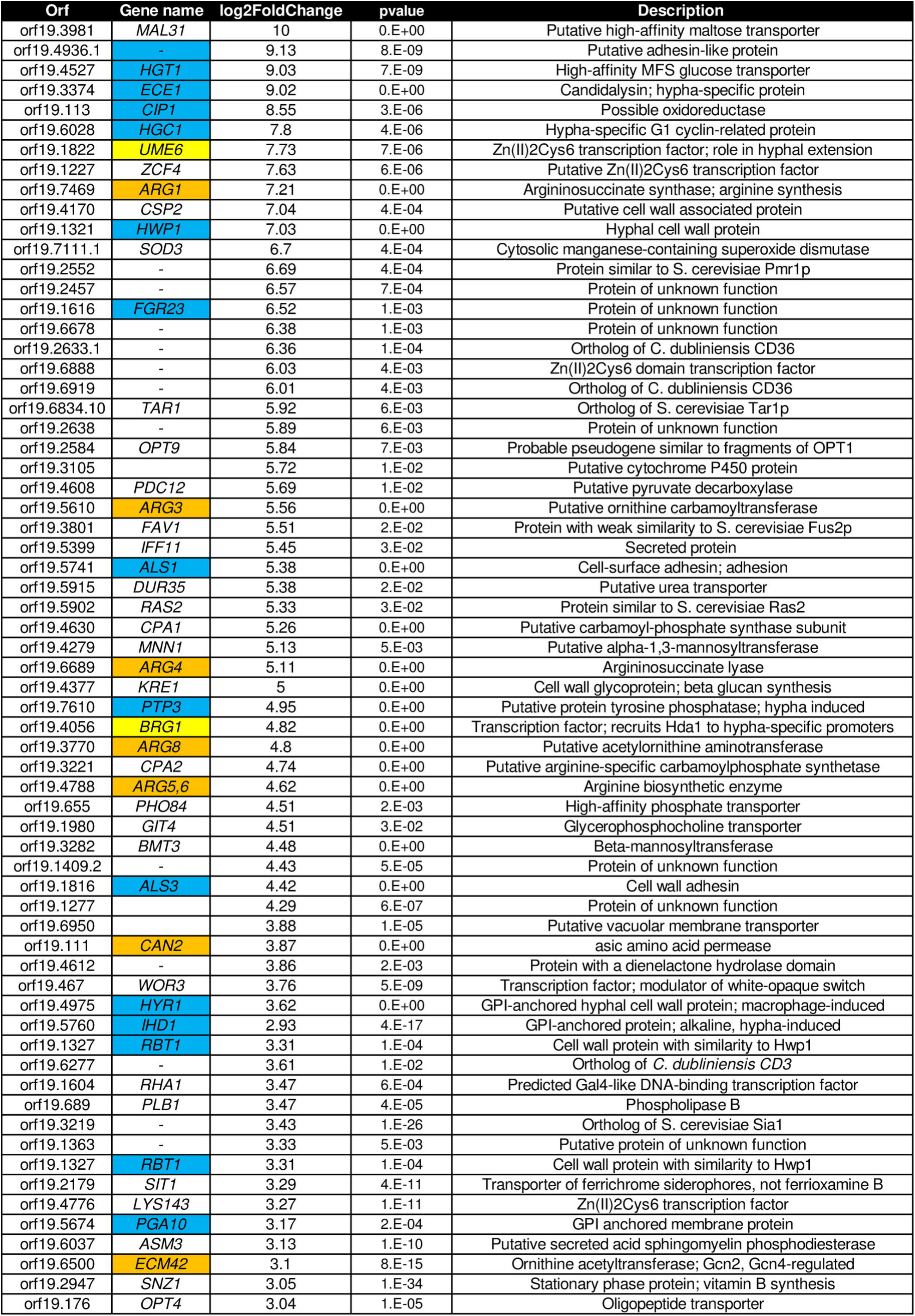
RNA sequence result of Rha1-GOF. Duplicated cultures of WT and RHA-GOF strains were grown overnight and processed for RNA seq. Upregulated classes of genes identified include genes involved in *Candida albicans* filament formation and arginine biosynthesis. Arginine biosynthesis related genes are shown in orange, filament related genes are in blue, and hyphal specific transcription factors are in yellow. The selected up-regulated genes are above a 3.5 fold cut-off with *P*-values <0.03.

### Loss of Rha1 function modulates hyphal development but not arginine prototrophy

Because activation of Rha1 leads to filamentation, we asked if the Rha1 function is required for normal hyphal development in response to hyphal inducing conditions. We inactivated both alleles of *RHA1* in the SC5314 background and assessed the phenotypic impact. The mutant cells were essentially unchanged in response to cell wall stresses and flocculated normally, but formed somewhat compromised biofilms (Figure 1A-E). We also assessed hyphal development in response to both serum induction and growth on Spider medium at 37°C. Colony morphology appeared unchanged under solid serum media hyphal inducing conditions (Figure 5A) but showed a distinct phenotype on Spider medium (figure 6A). There was a definite impact on cellular behaviour in response to either hyphal inducing condition in liquid medium. As shown in figure 5B, the null mutants were less filamentous under serum inducing conditions at 37°C at time 3.5 h with a defect in aspect ratio as compared to the wild type (Figure 5C). The filamentation defect was even more extreme for Spider induction conditions shown in figure 6B; here, the *rha1* null mutant strain was strongly compromised in hyphal development after 4 hours at 37°C. We assessed whether the *rha1* null strain was auxotrophic for arginine because activation of Rha1 also leads to the up-regulation of genes involved in arginine metabolism. We grew the strain in liquid SC-arg medium for 24 h at 30°C. In the absence of arginine supplementation; the mutant strain grew identically to the control strain (data not shown).

**Fig 5.**
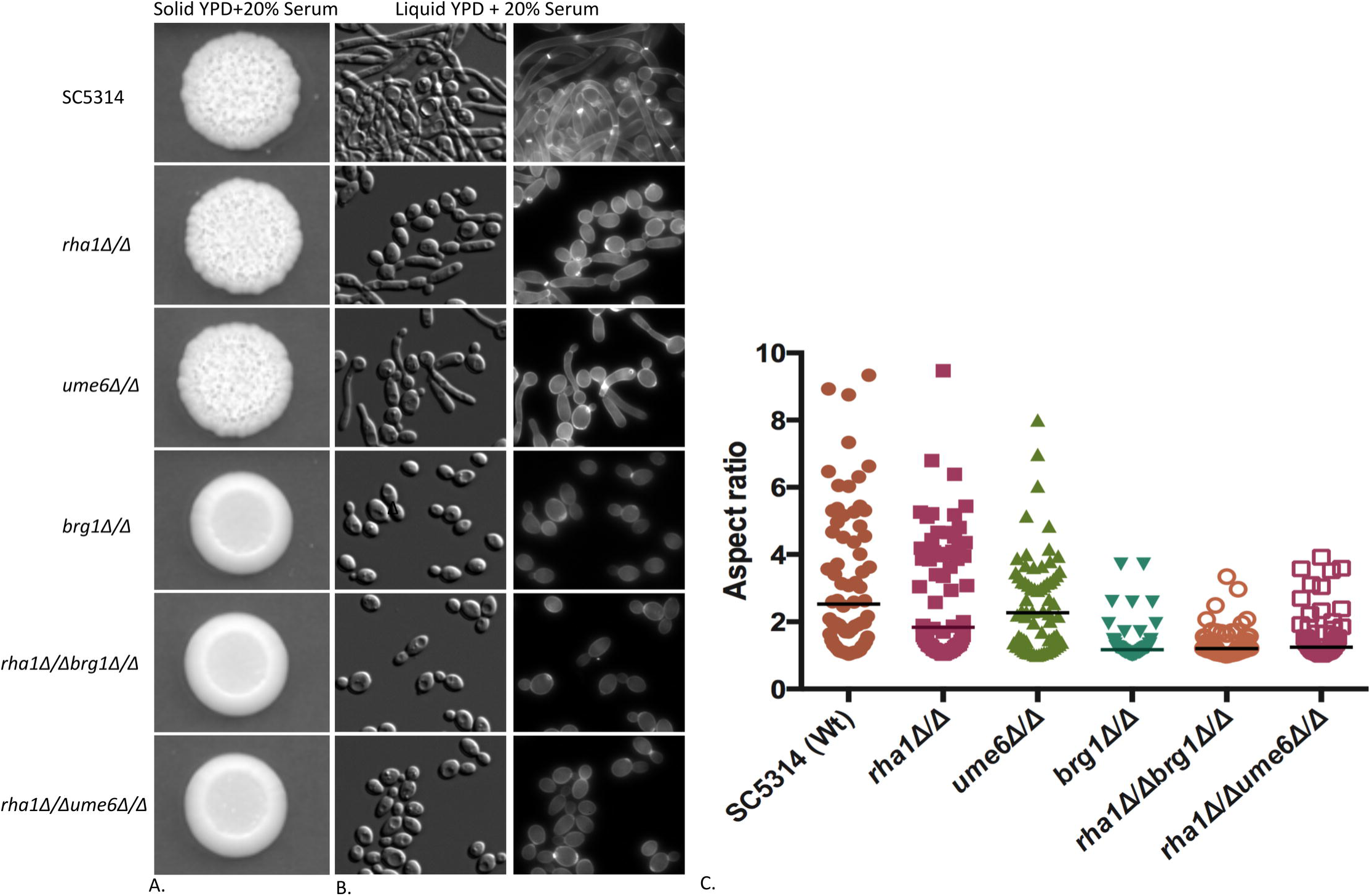
Deletion of *rha1* Δ/Δ and *rha1* Δ/Δ Ume6Δ/Δ causes variants to defect under serum stimuli. (A) The wrinkled colony morphology of *rha1* and *ume6* single mutants under 20% solid serum medium are shown after three days (B) The cellular morphology of the treated strains grown in liquid YPD supplemented with 20% serum and stained with CFW. (C). Aspect ratio analysis of the cellular morphology of strains with the single mutant of *brg1* and double mutants of *rha1Δ/Δ ume6Δ/Δ* grown under serum stimulation. Images were taken by DIC microscope; N≥70 cells

**Fig 6.**
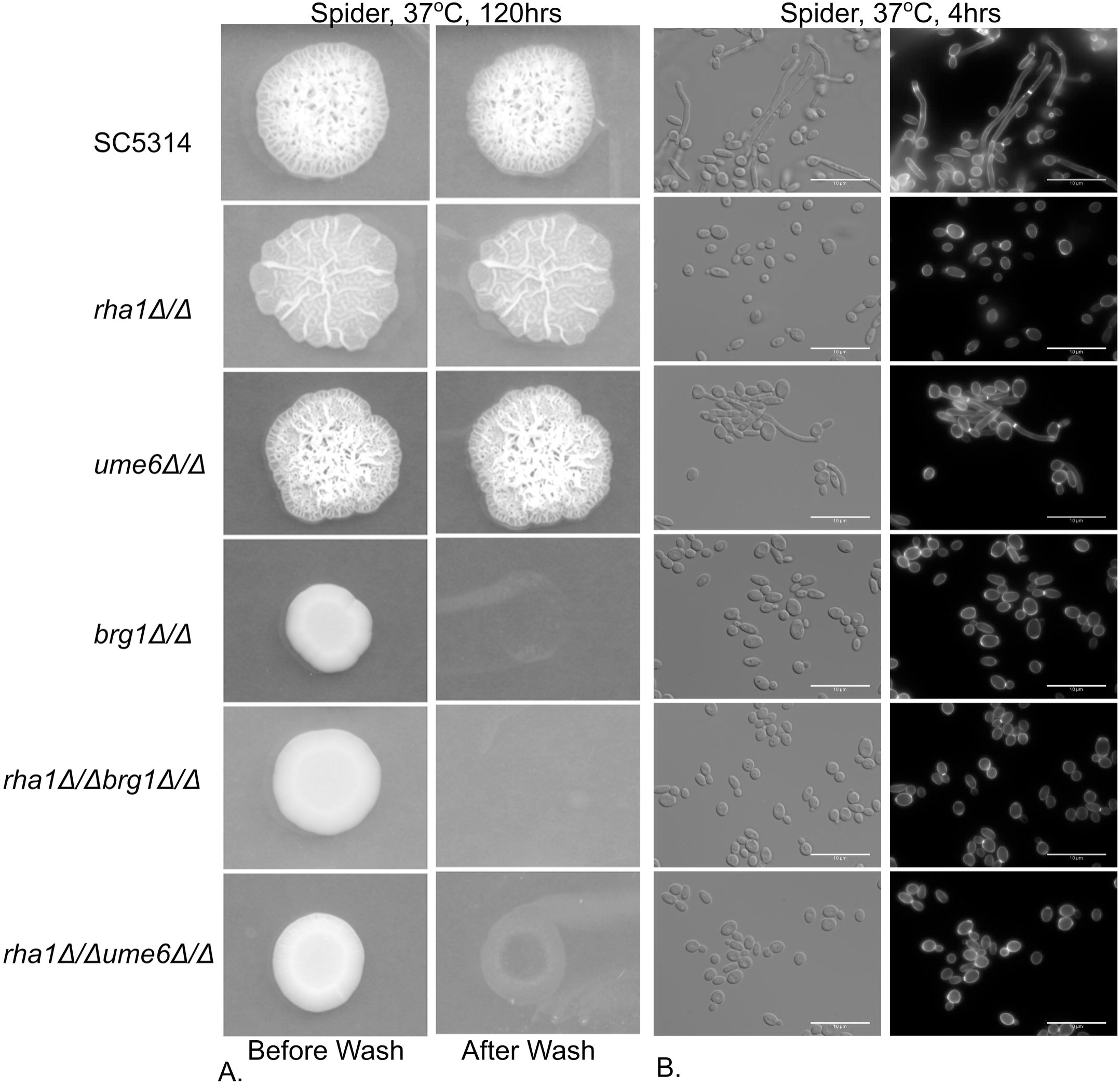
Deletion of Rha1 causes defects in hyphal formation on Spider medium. (A) The invasiveness of strains after growth on Spider medium for 120hrs was tested after washing colonies with a stream of water. (B) The cellular morphology of the indicated strains grown in the liquid Spider medium. The double mutants of *rha1 Δ/Δ Ume6Δ/Δ* were highly defective in hyphal formation when grown in Spider medium.

### Brg1 and Ume6 are required for the Rha1 hyperactive phenotype

The expression profiling evidence showed that the key hyphal-regulator-encoding genes *BRG1* and *UME6* were overexpressed in strains with hyperactive Rha1. We investigated whether the function of either gene was implicated in the filamentous phenotype generated by Rha1 hyperactivation under normal yeast growth conditions. We deleted *UME6* or *BRG1* in strains containing the activated Rha1 construct; as shown in Figure 7, we observed a severe loss of induced filamentation in the *brg1Δ/brg1Δ RHA1-GOF*, and a noticeable reduction in filamentation in the *ume6Δ/ume6Δ RHA1-GOF* strain.

**Fig 7.**
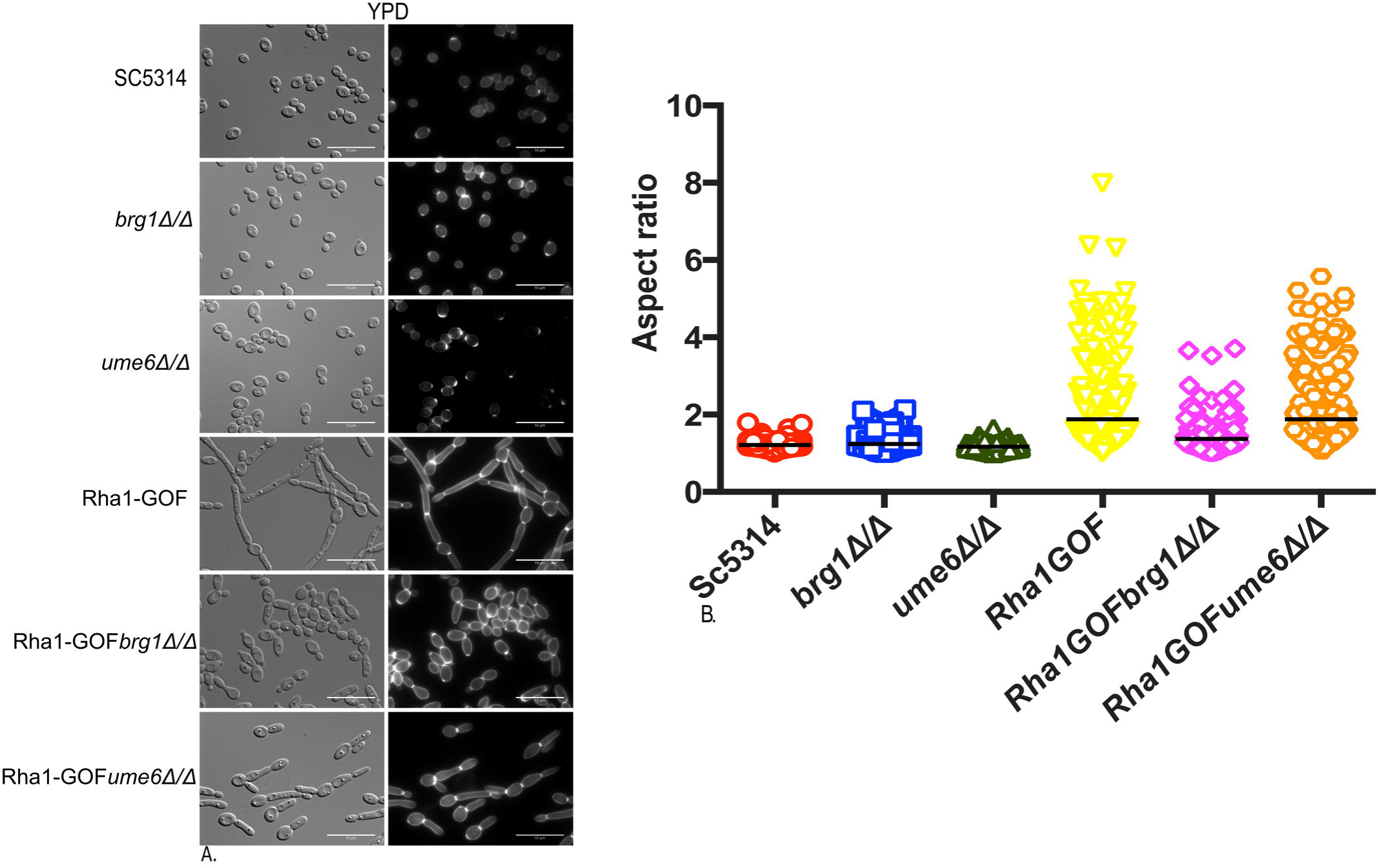
Deletion of Brg1 and Ume6 transcription factors causes defects in Rha1-GOF-induced morphology. (A) Cellular morphology of the indicated strains under yeast growth conditions and imaged with DIC optics (B) Aspect ratio of indicated cells N≥200; deletion of *brg1* modified the morphology of Rha1-GOF strains.

### Rha1 is part of a network of TFs controlling *C. albicans* filamentation

We further investigated the relationship between Rha1 and these other hyphal-control transcription factors identified in our profiling experiment. We first constructed double mutant combinations of the *RHA1* null with *UME6* and *BRG1* nulls to extend our examination of the effect of *RHA1* on hyphal development in response to environmental stimuli. As shown in Figure 5 and Figure 6, the *rha1 brg1* double mutant was not significantly different from the *brg1* null, which was by itself essentially non-hyphal in response to either serum or Spider inducing conditions. By contrast, however, the *rha1 ume6* double mutant was severely compromised in hyphal formation in response to either serum or Spider stimulation, even though the single mutants still showed significant morphological response to either hyphal inducing condition (Figures 5 and 6).

We also examined the effect of the Rha1 activated allele on the serum and Spider medium stimulation of hyphal development in the *UME6* and *BRG1* null mutants. As can be seen in figure 8, cells that were defective in hyphal formation in response to serum due to deletion of *UME6* or *BRG1* formed long filaments when the environmental stimulus was combined with the hyperactivated Rha1 construct. Serum treatment in the presence of the hyperactive Rha1 construct was able to stimulate some cellular elongation even in cells deleted for both *BRG1* and *UME6* (Figure 8). As well, strains with activated Rha1 and *brg1* or *ume6* deleted were somewhat filamentous on Spider medium (Figure 9).

**Fig 8.**
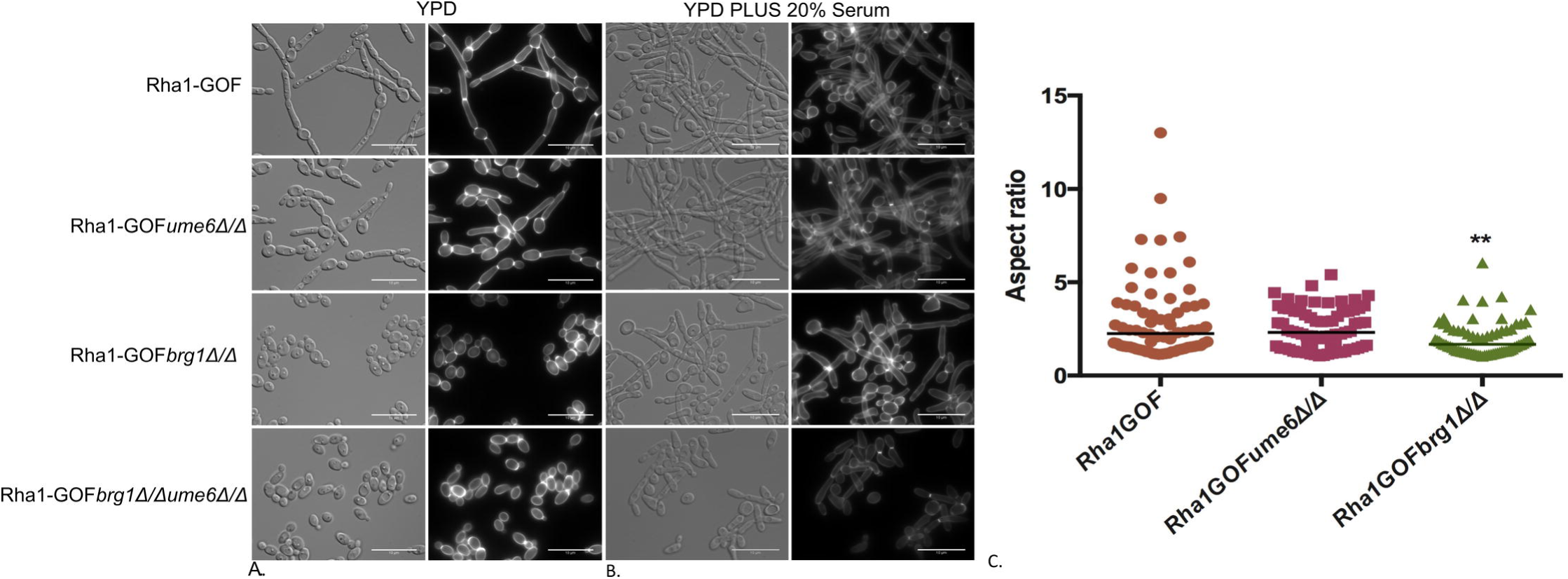
Strains expressing Rha1-GOF and supplemented with 20% serum rescue *brg1* and *ume6* single mutants for filament development. (A) Overnight cultures of strains grown in YPD plus 20% serum (B) The aspect ratio of cells after 3.5hrs in 20% serum and overexpression of Rha1. Supplemented with serum of Rha1-GOF could bypass the effect of single *brg1* and *ume6* mutants. Number of Cells N≥50 on the three different occasions.

**Fig 9.**
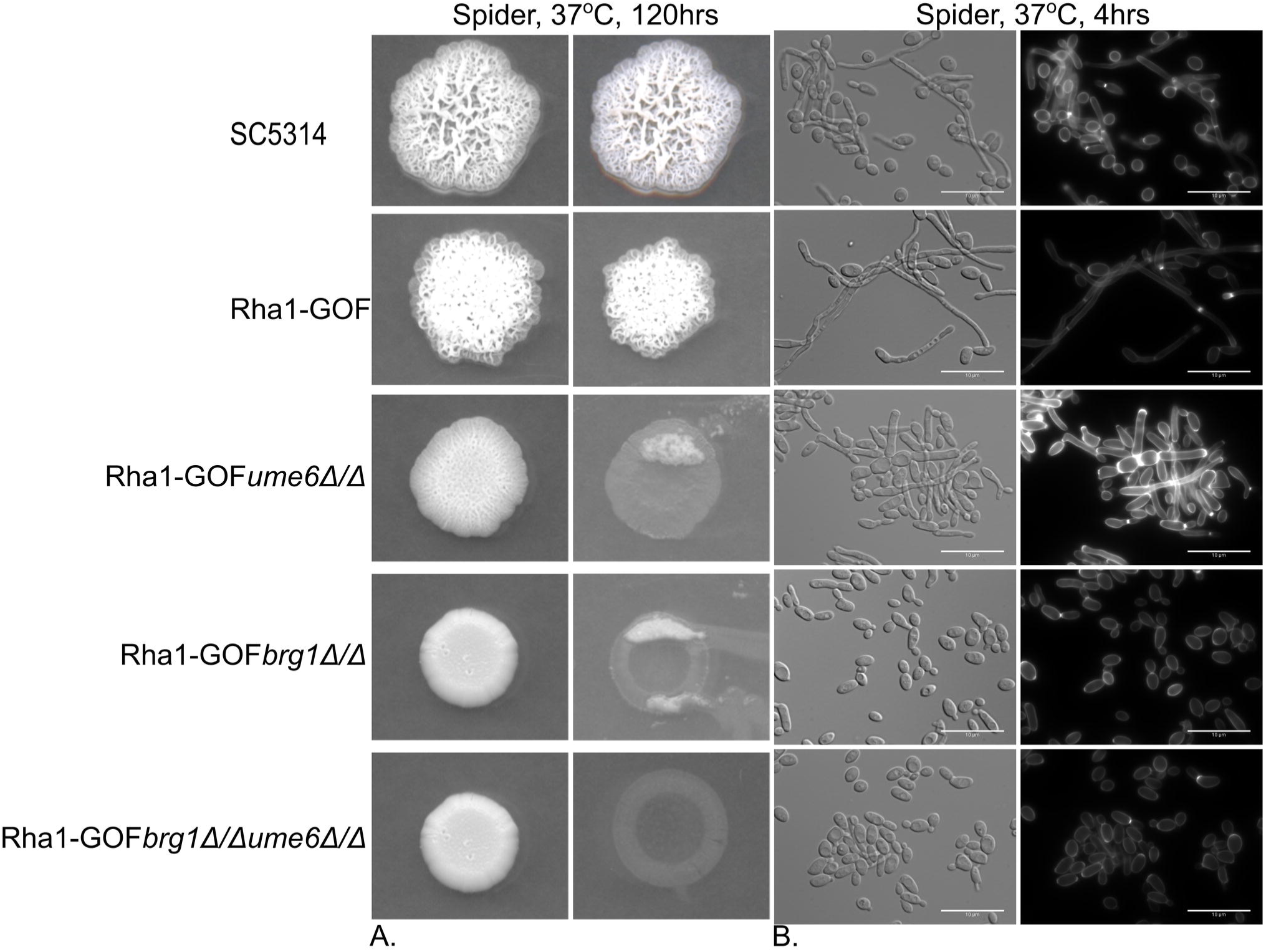
The media specificity of Rha1 for bypass of the filamentation defects of single *brg1* and *ume6* mutants. (A) Strains were spotted on Spider medium, and invasiveness assayed after being washed twice with 1xPBS and incubated 5 days. (B) Cellular morphology of strains grown in liquid Spider medium. Rha1 overexpression coupled with Spider medium showed some morphological effects on the *ume6* null strain but failed to bypass the consequences of *brg1Δ/Δ* or *brg1Δ/Δ um6Δ/Δ*.

## Discussion

One of the critical characteristics that allows *Candida albicans* to be a successful fungal pathogen is its capability of transitioning from a budding yeast to a filamentous form. *C. albicans* cells use this morphological plasticity to escape from human macrophages and to invade into deeper tissues during infection (11, 13, 25). Numerous studies have identified a variety of signaling pathways, with a complex interconnected network of transcription factors (TFs), required for regulation of hyphae associated genes (HAG) under different external stimuli (26). These signaling pathways include the mitogen-activated protein kinase (MAPK), the target <u>o</u>f <u>r</u>apamycin (<u>TOR</u>), the <u>r</u>egulation of <u>A</u>ce2 <u>m</u>orphogenesis (<u>RAM</u>), and the cAMP/RAS pathways (27).

The use of gene deletions to investigate the roles of individual members of such complex transcriptional regulatory circuits can be problematic in cases of functional redundancy (28). As an alternative, artificial gene activation can be a powerful strategy to uncover specific gene functions and to relate a phenotype to genotype for functional discovery in such interlinked systems (29). In this study, we use a variety of approaches, including genetic activation, to explore the molecular role of a new member of the hyphal control circuit, a zinc cluster family member we have termed Rha1 for regulator of hyphal activity. The Rha1 protein is a classic zinc cluster transcription factor; it is 989 amino acids long, with the zinc cluster DNA binding domain found from aa 16 to aa 47. Protein sequence alignments of CaRha1p reveal putative orthologs exclusively in the CTG clade of Saccharomycotina, whereas searches directed outside of the Candida clade show sequence similarity limited to the generally conserved zinc cluster DNA binding domain. The CTG group contains a large number of closely related pathogenic yeast, including *Candida dubliniensis, Candida parapsilosis*, *Candida tropicalis*, *Candida lusitaniae*, and *Candida guilliermondii* and thus Rha1 function may be associated with pathogenesis. Activation of Rha1 establishes that it functions in the hyphal development circuit; transcriptional profiling and analysis of genetic interactions suggest that its role in this circuit connects it to chromatin structure through the Nrg1 repressor and the Brg1 and Ume6 activators.

Regulation of transcription factors such as Brg1 and Ume6 has been connected to a central signaling pathway controlling hyphal development that requires the cAMP-dependent Protein Kinase (PKA). PKA is activated by a cAMP signal created through Ras1 and adenylyl cyclase (Cdc35/Cyr1). The action of this pathway ultimately results in the relief of Nrg1-mediated repression of hyphal associated genes (HAGs) (19, 30); overall cyclic AMP-dependent PKA controls chromatin conformation by tipping the state at the promoters of the HAGs from Nrg1-mediated repression to Brg1-mediated activation. This transition also involves a subunit of the histone chaperone complex (Hir1), which has been shown to be involved in the downregulation of Nrg1 and initiation of hyphal development (31), and the hyphal elongation transcription factor Ume6, which functions during the hyphal maintenance phase (26).

Strains defective in Brg1 are unable to initiate hyphal development and remained locked in the yeast state in response to activating signals like serum (23, 32). Strains defective in Ume6 can initiate but not maintain induced hyphal development, and thus generate short germ tubes under inducing conditions (21, 22). However, both the *brg1* null and the *ume6* null were bypassed in serum-stimulated cells that contained a hyperactive allele of Rha1; hyphal development in response to serum proceeded efficiently, suggesting Rha1 hyperactivity can compensate for the loss of either Brg1 or Ume6. However, in the presence of hyperactivated Rha1, serum treatment of the *brg1 ume6* double mutant triggered polarized growth but not extensive filaments, showing that Rha1 activation cannot altogether bypass the entire Brg1/Ume6 circuit. Intriguingly, on solid medium hyperactive Rha1 did not trigger bypass of the *brg1* mutant in the presence of serum. Previous studies have established different genetic and transcriptional programs during hyphal development in cells responding to growth in either liquid or solid media (33).

In addition to bypassing blocks in serum-stimulated hyphal development in *brg1* and *ume6* mutations, we found that hyperactive Rha1 directs constitutive hyphal growth, creates an invasive phenotype in the absence of external hyphal inducing signals, and stimulates biofilm formation. As well, the activated Rha1 strains are hypersensitive to cell wall stressors such as Caspofungin and Congo Red, suggesting Rha1 hyperactivity may influence cell wall composition. Consistent with the phenotypic observations, transcriptional profiling of the hyperactivated Rha1 strain in the absence of external signals for hyphal development reveals induction of a variety of hyphal-specific genes like *ECE1* and *HWP1*, as well as genes for the transcription factors Brg1 and Ume6, and also shows downregulation of the gene for the hyphal repressor Nrg1. The induction of filamentation caused by Rha1 activation in the absence of external signals depends entirely on Brg1, while loss of Ume6 reduces, but does not eliminate, Rha1-hyperactivity-induced filamentation. The deletion of both Brg1 and Ume6 together completely blocks the filamentation caused by hyperactive Rha1 under yeast growth conditions. Previous work (34) has shown that *RHA1* is repressed in yeast cells by the Nrg1p–Tup1p transcriptional repressor pair, although further study is required to establish if this repression is direct. Overall then, hyperactivation of Rha1 in the presence or absence of serum stimulation establishes an active link between Rha1 function and the Nrg1-Brg1 switch regulating chromatin structure at HAG promoters and the initiation of the hyphal developmental program.

The deletion of *RHA1* also provides evidence for a link between Rha1 and the Nrg1/Brg1-Ume6 circuit. The deletion mutants show limited defects in hyphal development, as filamentation is moderately delayed in serum-stimulated cells, and more significantly delayed in Spider medium induced cells. However, the *rha1 ume6* double mutant is utterly defective in the serum of Spider induced hyphal development, although either mutant alone was capable of initiating hyphal formation. This suggests some redundancy in the roles of Rha1 and Ume6 in the hyphal developmental program.

Interestingly, Rha1 may function in more than just the yeast to hyphal transition. Genes involved in arginine biosynthesis are also up-regulated in cells with a hyperactivated Rha1. Previous work has suggested a linkage between hyphal development and arginine biosynthesis (35); Rha1 may thus play a role in connecting these two cellular processes. Overall our study provides insight into the mechanism of hyphal development controlled by the Nrg1/Brg1 switch. Under hyphal inducing conditions, Rha1 functions to facilitate both downregulation of the Nrg1 repressor and upregulation (directly or indirectly) of the Brg1 and Ume6 transcription factors required for initiation and maintenance of HAGs. This switches the cell from a repressed chromatin configuration at the HAGs controlled by Nrg1 to the activated state controlled by Brg1, and thus Rha1 represents an important new regulator of the yeast to hyphal transition critical for *C. albicans* pathogenicity.

## Materials and Methods

### Strains, media and growth conditions

All *Candida albicans* strains, oligonucleotides, and plasmids used in this study are listed in tables S1 and S2. For long term storage, cells were kept at -80°C in 25% glycerol supplemented YPD. The strains were routinely cultured in liquid YPD (10 g yeast extract, 20 g peptone, 20 g glucose per liter, Uridine (50 μg ml−1), 2% w/v Bacto-agar for solid medium) at 30°C in a shaking incubator. The *C albicans* transformations were plated on YPD agar plates containing 200 µg/ml nourseothricin acetyltransferase gene (Nat1) [Werner Bioagents, Jena, Germany) or 500-600 µg/ml Hygromycin B (HygB) (Bioshop Inc, Canada), for 48 h at 30°C]

### *C. albicans* mutant strains

All *C. albicans* mutants were constructed in the wild type strain SC5341. The lithium acetate method of transformation (36) was used with the modification of growing transformants overnight in liquid YPD at room temperature after removing the lithium acetate-PEG. A CRISPR/Cas9 system (37) was performed to construct an RHA1 deletion strain. The sgRNA of *RHA1* was formed by annealing primers RHA1-sg-F and RHA1-sg-R and cloning the fragment into the BsmBI site of pV1093 to make plasmid pV1093-Rha1-sgRNA. A repair DNA was amplified using Rha1-Rep-F, and Rha1-Rep-R primers to PCR amplify fragment containing in-frame stop codons with a disrupted PAM region and introduction of a restriction enzyme cleavage site for the confirmation of transformants with the correct insertions. The repair DNA template and linearized plasmid pV1093-Rha1-sgRNA was transformed then the correct deletion was verified by PCR. We used a transient CRISPR/CAS9 system (38) to delete *C. albicans BRG1* and *UME6* in the wild type SC5314 and Rha1-GOF background strains. A Cas9 gene was amplified using a pV1093 plasmid and P7 and P8 standard primers. The final sgRNA fragment was amplified using P5 and P6 primers from the product of two separate PCR reactions, including the sgRNA sequence. The HygB repair template was amplified from a HyB plasmid as a selection marker to create homozygous null mutants *brg1 Δ/Δ* and *ume6 Δ/Δ*. correct deletions were confirmed using primers internal to the HygB markers in combination with primers for the upstream regions of *UME6* or *BRG1* and by using primers internal to the *BRG1* and *UME6* ORFs. The double mutant strains *rha1 /rha1 ume6/ume6* and *rha1/rha1 brg1/brg1* were constructed in the homozygous deletion of *rha1 /rha1* Nat^R^ strain background.

### Filamentation assays

The environmental filament induction assays were performed by growing the selected strains overnight in liquid YPD at 30°C, 220rpm. The next day, the cells were washed twice with 1X PBS and resuspended at OD_600_ = 0.1 into a fresh liquid Spider (Liu et al., 1994), YPD, or YPD+20% serum (Fetal Bovine serum; Sigma) then incubated at 37°C for 3.5-4 hours. For solid medium, 5 µl from an OD_600_ = 0.1 culture was spotted onto the YPD plus 20% serum plate and incubated at 37°C for 3 days.

### Biofilm assays

Single biological replicates from the selected strains were grown overnight in 5ml liquid YPD at 30°C on a shaker at 220-rpm. In the morning, after washing the cells twice with 1XPBS, the cell densities were adjusted to an OD600=0.5 in 200 μl Spider medium and then were added to a 96 well assay for 24 hrs. Each well was washed twice with 200 μl of PBS, and the level of biofilm formation was determined by using the crystal violet assay, as reported previously (39).

### Stress assays

For stress assays, single colonies from strains Sc5314 (control), *RHA1-*GOF and the *rha1* null were grown overnight in YPD at 30°C and resuspended in 1XPBS to a cell density of 1.5×10 ^7^ cells/ml at 600nm, then were diluted in 10-fold stages from 10^6^ to 10^2^ in 1X PBS. These tenfold serial dilutions of each strain were spotted onto YPD plates comprising the indicated compounds Caspofungin (0.75 μg/ml), Congo red (200 μg/ml), fluconazole (10 μg/ml), hydrogen peroxide (5 mM), MMS (0.02% v/v), 10mM Fe (II) SO4, 20 mM Fe (III) Cl, and 5mM CuSO4, 0.4m Cacl2, the osmotic stressor 1M sorbitol, Menadione 0.15mM, HU 38mM, PH10, Glycerol 250mM, Hygromycin B 100Mg/ml. Plates were incubated for 3 days at 30°C.

### Microscopy and imaging

Single colonies of *C. albicans* strains were inoculated in 5 mL of liquid YPD for overnight, at 30°C, and shaken at 220-rpm. The cells were then washed twice with 1xPBS and imaged using differential interference contrast (DIC). For the hyphal induction images, cells were washed twice with 1XPBS, followed by the addition of Calcofluor staining (2 μg/ml) for 20min. DIC examined cells at 63× and ×100 magnification using a Leica DM 6000 microscope. For quantification of elongation factor, cells were stained with Calcofluor white (Sigma, USA.), then imaged using a Leica DM6000 microscope with both DIC optics and appropriate fluorescent filters, using a 100x (N.A. 1.3), 60x (N.A. 1.4) or 40x (N.A. 0.75) objective lens. After capture, Calcofluor white images were presented to a Region Convolutional Neural Net (40) trained to recognize yeast cells, resulting in a binary mask that represents the outline of most yeast cells in the image; these masks were verified by a trained human observer, who could discard inappropriate masks that did not correlate well with merged DIC and Calcofluor white images. The remaining masks were measured in FIJI (NIH, Bethesda), using the Shape Descriptors option to extract the aspect ratio of each cell.

### Invasion assays

Single colonies of the selected strains were grown overnight in 5 mL YPD at 30 °C 220-rpm shaker. Then the cells were washed twice with 1x PBS and diluted to an optical density of OD_600_=0.1. 5 µL from the adjusted cell density were spotted on YPD, 1 day at 30 °C or Spider agar at 37 °C, 5 days. The spots on the spider plates were then washed with sterile water for 15 seconds. Plates were scanned before and after washing using an Epson Perfection V500 Photo color scanner.

### RNA isolation and RNA-seq experiment

Two biological replicates of both SC1604GAD1A and the Wild-type SC5314 strains were diluted to OD600=0.1 in YPD and inoculated to reach OD600 =0.8-1. Total RNA was extracted using Qiagen RNeasy plus Minikit, then RNA quality and quantity were determined using an Agilent bioanalyzer. Paired-end Illumina Illumina miSEQ sequencing of extracted RNA samples was carried out at the Quebec Genome Innovation Center located at McGill University. Raw reads were pre-processed with the sequence-grooming tool cutadapt version 0.4.1 (MARTIN 2011) with the following quality trimming and filtering parameters (‘--phred33 --length 36 -q 5 --stringency 1 -e 0.1’). Each set of paired-end reads was mapped against the C. albicans SC5314 haplotype A; version A22 downloaded from the Candida Genome Database (CGD) (http://www.candidagenome.org/) using HISAT2 version 2.0.4. SAMtools was then used to sort and convert SAM files. The read alignments and SC5314 genome annotation were provided as input into 13 StringTie v1.3.3 (PERTEA et al. 2015), which returned gene abundances for each sample. Raw and processed data have been deposited in NCBI’s Gene Expression Omnibus (EDGAR et al. 2002).

## Acknowledgments

We want to acknowledge funding support from the Tier I Canada Research Chair program and from CIHR grant MOP42516 to M.W., a Concordia University Faculty of Arts and Science Graduate Fellowship. We want to thank Dr. Joachim Morschhäuser for the Rha1-Gof*um6*Δ/Δ deletion strain. We are grateful to Dr. Chris Law of the Centre for Microscopy and Cellular Imaging for assistance with microscopy and the elongation factor assay.

## Figure Legends

**Fig 1S. The effect of different stresses on strains expressing Rha1-GOF**. Overnight culture of Sc5314 and Rha1-GOF were spotted under the indicated stress conditions. Rha1-GOF generally grow normally under the different stresses

