## Supplementary figures and images for "The zinc cluster transcription factor Rha1 is a positive filamentation regulator in *Candida albicans*"

### Supplemental Figure 1

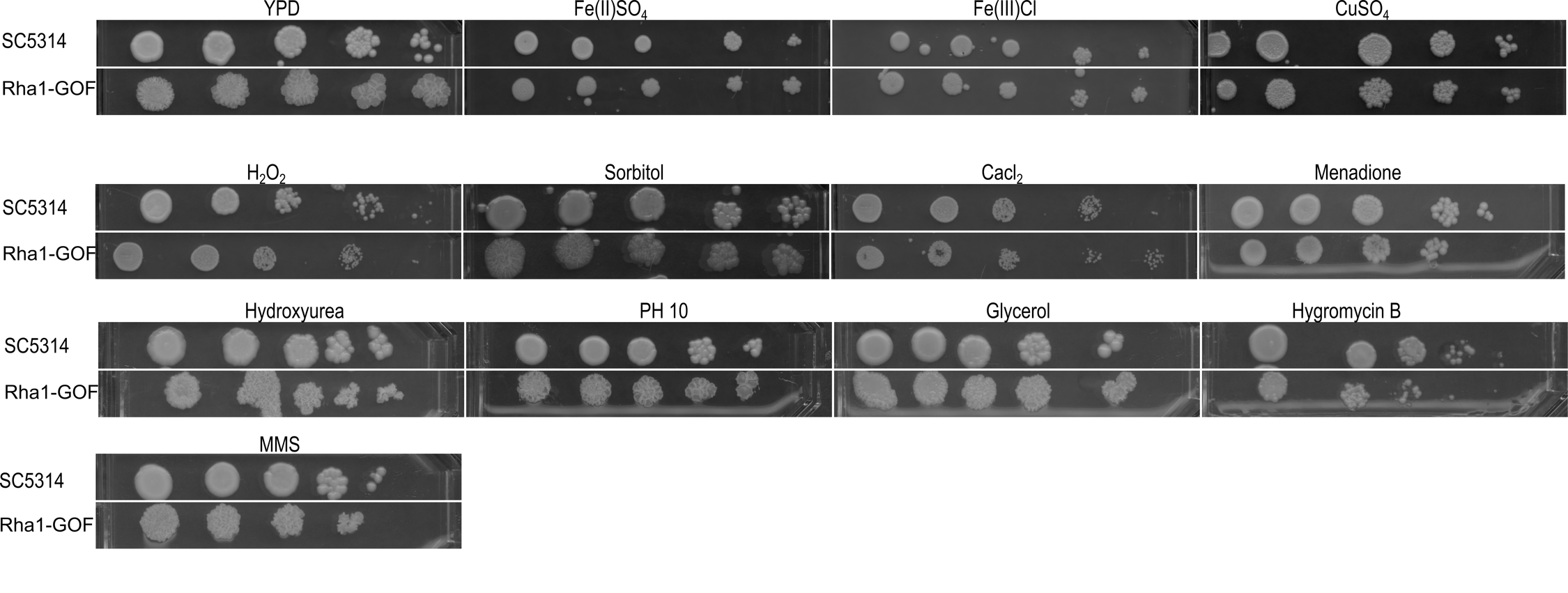
